# Effects of coral colony morphology on turbulent flow dynamics

**DOI:** 10.1101/839902

**Authors:** Md Monir Hossain, Anne E. Staples

## Abstract

Local flow dynamics play a central role in physiological processes like respiration and nutrient uptake in coral reefs. Despite the importance of corals as hosts to a quarter of all marine life, and the pervasive threats currently facing corals, little is known about the detailed hydrodynamics of branching coral colonies. Here, in order to investigate the effects of the colony branch density and surface roughness on the local flow field, three-dimensional simulations were performed using immersed boundary, large-eddy simulations for four different colony geometries under low and high unidirectional oncoming flow conditions. The first two colonies were from the *Pocillopora* genus, one with a densely branched geometry, and one with a comparatively loosely branched geometry. The second pair of colony geometries were derived from a scan of a single *Montipora capitata* colony, one with the verrucae covering the surface intact, and one with the verrucae removed. We found that the mean velocity profiles in the densely branched colony changed substantially in the middle of the colony, becoming significantly reduced at middle heights where flow penetration was poor, while the mean velocity profiles in the loosely branched colony remained similar in character from the front to the back of the colony, with no middle-range velocity deficit appearing at the center of the colony. When comparing the turbulent flow statistics at the surface of the rough and smooth *M. capitata* colonies, we found higher Reynolds stress components for the smooth colony, indicating higher rates of mixing and transport compared to the rough colony, which must sacrifice mixing and transport efficiency in order to maintain its surface integrity in its natural high-flow environment. These results suggest that the densely branched, roughly patterned corals found in high flow areas may be more resistant not only to breakage, but also to flow penetration.

## Introduction

A new and important direction in understanding the coupled dynamics of corals and their hydrodynamic environments seeks to measure and model the flow inside coral reefs and individual colonies. But the task is challenging due to the existence of a wide range of flow scales within different boundary layers around a coral reef [1, 2]. Large-scale motions transport planktonic food to the reef, while diffusion [3, 4] takes place at a much smaller scale at the coral surface. For the smaller scale processes, experimental studies [5, 6] show that growth direction, dimensions, and sparsity of branches depend on the flow profile inside the coral [7–9]. Similarly, the velocity profile controls the nutrient distribution [7, 10–13]. and physiological processes like photosynthesis and respiration [14–17] near the coral surface. Flow motion also controls the thermal micro-environment [18, 19] at the coral surface, which is extremely important for incidents like coral bleaching. Both the transfer of nutrients from the water column to the coral and the transfer of byproducts from the coral to the water column depend on the details of the flow through and around the coral [20, 21]. It is clear from the discussion that all of these physical parameters are directly related to the flow conditions in and around the coral geometry. To obtain a clear picture of this interdependent relationship, the authors tried to find answers to a few basic questions, including: How does the flow profile change within the coral canopy?, Is it possible to quantify these variations? and What are the effects of the internal structure of coral on the surrounding fluid?

Though numerous studies have been performed on coral reefs, these analyses lack a detailed study of flow inside a single colony. Most of these studies were performed either above the coral canopy or on a complete reef [22, 23] without considering the arbitrary shape of geometry. One of the reasons for the lack of such critical study might be the complexity of measuring the flow parameters in close proximity to the coral’s surface. From an experimental point of view, examining flow inside of coral has proven difficult due to a lack of optical and acoustic access to the interior. Though few studies were performed on single corals, data obtained at several points were not sufficient to comprehend the complete flow profile inside of the coral. For past canopy flow studies, the models were composed of homogeneous, simplified elements and double averaged Navier-Stokes equations [24] were used to simplify the analysis, which led to modeling of a new unresolved dispersive stress [25, 26]. However, the analysis did not consider the heterogeneity of the structure, and the double-averaging procedure curtails the variation of flow at the inside of the coral. So, to understand the variation of the flow profile within a coral colony, it is necessary to resolve the three-dimensional velocity component on the inside of real coral with natural geometry, which is the focus of current study.

To understand the flow dynamics on the inside of the coral reef, Lowe *et al.* [27] tried to capture the velocity profile and shear stress by modeling the coral as an array of cylinders. This model was also used to estimate the mass transfer from the surface of the coral, but these experimental analyses on simplified structures were not sufficient to depict real flow scenarios through the complex branching patterns of coral. Bilger and Atkinson [28] used engineering mass transfer [29] correlations to predict rates of phosphate uptake by coral colonies and assumed the top of coral colonies as a rough boundary. Though this analysis is useful in describing a boundary profile for a large coral reef, it does not make clear the impacts of the individual coral colony on the flow structure. Recently, Chang *et al.* [30] have used magnetic resonance velocimetry [31] to reveal the internal flow profile of corals for two different branching patterns. Though the results provided an excellent visualization of the flow field inside the coral, the study did not provide sufficient information regarding the change in the velocity profile along the length of the coral. Additionally, repeating such experiments on diverse coral structures seems a tedious task. To solve these problems, numerical simulations are an important tool.

For numerical simulations, complexity arises during capture of the arbitrary boundaries of coral geometry. Specifically, meshing around the arbitrary branches of coral is a challenging task. For body-fitted mesh, even the simplest geometries require efficient meshing algorithms to reduce the computational cost. To overcome this issue, numerical experiments in previous studies were performed by considering coral as an isotropic, porous medium, but questions emerged due to non-consideration of coral’s natural morphology [32]. In addition, Kaandorp *et al.* used the Lattice Boltzmann method to understand the effects of flow on coral structure and simulated flow patterns around the coral surface [33,34] at low Reynolds numbers (154 to 3,840), due to stability issues. Similarly, Chindapol *et al.* [35] used COMSOL Multiphysics to analyze the effects of flow on coral growth only in a laminar framework. Chang *et al.* [36] were the first (according to the authors’ knowledge) to introduce immersed boundary method with large-eddy simulation on real coral geometry while computing the velocity profile at the inside of the coral, but the analysis was mainly focused on local shear and mass transfer on the coral structure. In a recent paper, Hossain and Staples *et al.* [37] computed the flow profile on the inside of coral, and showed that the mass transfer remains almost constant throughout the coral even when the velocity drops substantially inside it. However, the analysis did not provide sufficient information regarding variation in the mean velocity profile with the change in branching pattern, which will be discussed here in detail for two different *Pocillopora* branching structures.

In addition, it is important to understand the effects of natural flow conditions on coral morphology. To flourish in a changing environment, coral modifies its structure to adjust to the surrounding hydrodynamics. It has been found that under sheltered conditions, *Pocillopora damicornis* and *Seriatopora hystrix*, transformed from dense-branched to thin, larger-branched spacing coral due to the change in flow conditions [38]. Kaandorp and Kubler showed similar transformations for *Madracis mirabilis* [39]. Even a small modification can affect the boundary layer and turbulent stress developed at the coral surface and can change the mass transfer rate [40, 41]. These variations in turbulent flow profiles illustrate the survival mechanism for individual coral in different flow conditions. One example of such adaptability is the survival of loosely branched *Montipora capitata* in a strong flow environment. Normally, loosely branched coral grows in a quiet flow environment, but *Montipora capitata*, with verrucae on its surface, is found in highly turbulent flow conditions near Hawaii. Naturally, questions arise regarding the endurance of such loosely branched coral in a strong flow environment and the impact of the verrucae on the surrounding fluid. To understand these morphological adaptations, it is essential to comprehend and compare the formation of boundary layers and turbulent stresses on these altered coral structures of *Montipora capitata* with and without verrucae. Though there were numerous studies performed on coral reefs, capturing the morphological change with flow conditions is rare in the literature and is a challenging task. Here, a detailed comparative flow analysis was performed on *Montipora capitata* with and without verrucae to describe the adjustment of these living structures in strong and weak flow conditions.

Based on the discussion above, the current manuscript can be divided into two different sections. In the first segments, three-dimensional velocity components and mean velocity profile were computed at the interior of two *Pocillopora* corals with different branching patterns for the same incoming flow. In the second part of the manuscript, a comparative study was performed between velocity profiles and Reynolds stress developed on *Montipora capitata* with and without verrucae in quiet and strong flow conditions. Then, the results of the computations were analyzed to understand the development of velocity profiles and turbulent stress on the slightly altered coral structures for the same incoming flow.

## Materials and methods

### Coral species and geometries

Stereolithography (STL) files of the coral geometries used in the current studies are shown in Fig 1. Between the two *Pocillopora* coral, the densely branched *P. meandrina* was obtained from computed tomography (CT) of a real coral skeleton from Professor Uri Shavit’s laboratory in the Department of Civil and Environmental Engineering at the Technion - Israel Institute of Technology. *P. meandrina*, generally known as cauliflower coral, is common in the East Pacific and the Indo-West Pacific, and is mainly exposed to the reef front. The stream-wise length, width and height of *P. meandrina* used in the current simulation were 0.172 m, 0.172 m and 0.111 m respectively. In contrast, branches are distributed more sparsely in *P. eydouxi*. This coral is common throughout the Indo-West Pacific and occurs in most reef environments, especially at reef fronts where the currents are strong. The length, width and height of *P. eydouxi* used in the simulation were 0.12 m, 0.11 m and 0.085 m respectively. Though the genus was similar, there were substantial structural differences between these two coral morphologies. The ratio of volume to surface area for *P. meandrina* and *P. eydouxi* were 2.0 cm and 0.4 cm respectively. This parameter indicates the compactness of branches inside the coral colony, and the results showed that *P. meandrina* was almost five times more compact than *P. eydouxi*. The ratio of maximum projected area in flow direction to surface area were 0.0754 and 0.0556 for *P. meandrina* and *P. eydouxi* respectively, which represented a higher projected surface area for *P. meandrina*. To understand their structural differences more clearly, variations of branch diameter and cross-sectional area were plotted in Fig 1 with respect to height for both *Pocillopora* corals.

**Fig 1.**
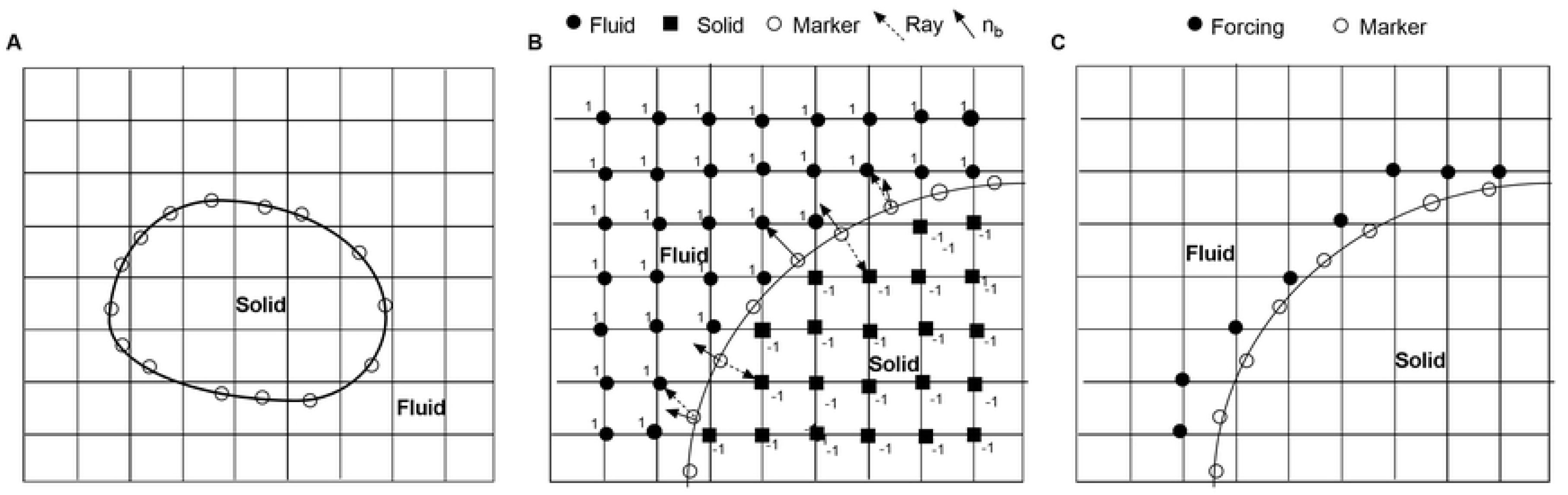
*Pocillopora* geometries used in simulations. (A, B) Variation of cross-sectional area through a horizontal plane and mean branch diameter versus height for *Pocillopora meandrina* and *Pocillopora eydouxi*, respectively. (C) Rendered stereolithography file of the *Pocillopora meandrina* colony used in the simulations and a *P. meandrina* colony in its natural environment on the ocean floor. (Brocken Inaglory, *Pocillopora meandrina* with a resident fish, Wikimedia Commons (2008); used in accordance with the Creative Commons Attribution (CC BY-SA 4.0) license [43]). (D) Rendered stereolithography file of the *Pocillopora eydouxi* colony used, and a *P. eydouxi* colony in its natural environment. (Paul Asman and Jill Lenoble, Black-axil and bluegreen chromis, *Chromis atripectoralis* and *C. viridis*, Wikimedia Commons (2012); used in accordance with the Creative Commons Attribution (CC BY-SA 2.0) license [44]).

To comprehend the effects of verrucae on the surrounding fluid, two coral geometries with and without verrucae were used for computation (Fig 8). Both coral geometries originated from the same STL file of *Montipora capitata*. Fig 8 (A) shows *M. capitata* without verrucae where the verrucae from the coral surface were completely removed while keeping the other parameters the same. The dimensions of both coral were similar. The length, width and height of the corals were 0.118 m, 0.114 m and 0.1 m respectively. The cone-shaped verrucae had a base diameter of approximately 3 mm and a length of 3–4 mm. *M. capitata* is usually found in highly turbulent flow conditions in the tropical north and central Pacific Ocean near Hawaii at approximately 20 m depth. All corals used in the current study had low living-tissue biomass, so using skeletons instead of coral with bio-tissue had a minimal effect on flow dynamics.

### Numerical methodology

The authors were looking for a method that could be implemented easily for complex geometries while minimizing computational cost. Immersed boundary method (IBM) is suitable for such analysis where the computation is performed on a Cartesian grid and the interface is tracked by identifying solid and fluid elements in the flow domain. For the current analysis, IBM was implemented in a large eddy simulation (LES) framework for a turbulent flow field where the largest scales of motion were calculated and the effects of the small scales were modeled. A top-hat filter was applied implicitly to the Navier-Stokes equations by the finite-difference operators and the resulting filtered equations are as follows:

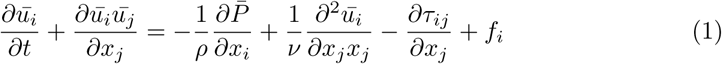

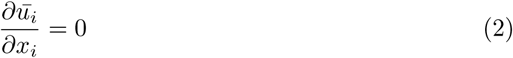

where 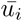 represents large-scale velocity vector, *P* represents pressure, *ρ* represents density, *ν* represents kinematic viscosity and *f*_*i*_ represents the external body force used to implement boundary conditions for arbitrary objects with a non-conforming grid. Here, the overbar denotes a filtered variable.

The spatial filtering operation on the Navier-Stokes equations produces the subgrid scale stress term,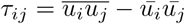, which needs a closure model to solve the system of equations (1) and (2). The dynamics eddy-viscosity model used in this study is of the following form:

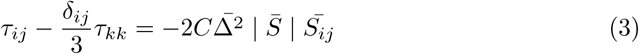

where 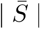 is the magnitude of the large-scale strain rate tensor, and 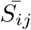 is strain tensor and 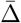 is the filter size, which can be defined as:

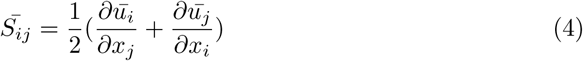

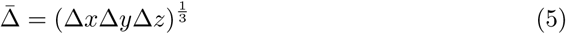

Here, ∆*x*, ∆*y* and ∆*z* are the local grid size, and C is the user-specified value for the Smagorinsky eddy viscosity model, where the usual value of C ranges from 0.1 to 0.2. However, the model does not behave properly near solid boundaries. So, in the computation analysis, the dynamic modeling procedure was used to compute the value of C from the information of the resolved scale [45]. All of the three-dimensional simulations on STL files of coral colonies were performed using the IB method implemented in the LES framework using a code developed by Balaras *et al.* The complete details of the numerical method can be found in Balaras *et al.* [46].

A standard second-order central-difference scheme on a staggered grid is used in the present study. The pressure and scalar variables are located at the center of the grid cell, and velocity components are located at the cell-face center. The solution of equations (1) and (2) is obtained by a second-order projection method. Here, time was advanced explicitly by using an Adams–Bashforth scheme. An explicit algorithm has been used to simplify the implementation of boundary conditions near a complex boundary. In a two-step time-splitting method, first a provisional value of the velocity field, which is not divergence free, can be obtained and used for updating the velocity. The intermediate velocity, 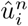 evaluated at n step can be written as follows:

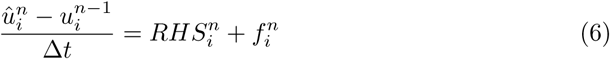

where RHS contains all of the terms in the right side of the momentum equation. After solving the Poisson equation, the final velocity, 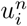, can be updated from the 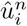. The external force, 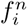, needs to be calculated first to implement the desired boundary condition at the boundary point where the boundary may or may not coincide with the grid points. To do this, it is necessary to identify solid, fluid and forcing points near the boundary where forcing functions are implemented in the simulation domain. In the current study, accurate interface tracking methodologies were used for complex coral geometry. An immersed solid boundary of an arbitrary shape is represented by a series of markers shown in Fig. 2. After defining the immersed interface as a series of marker particles, the relationship between these particles and the underlying Eulerian grid can be established by ray-tracing algorithm. The procedures for identifying solid and fluid points in the computational adopted for the current study can be found in detail in [46]. The procedure includes Finding of the closest grid points near each marker particle. Then from each marker particle, a ray 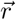 is shot to the grid point and a normal, 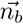, is also calculated on the marker points. If 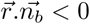, then this grid point is inside the solid; otherwise, it is outside of the solid. The forcing points are the closest grid points near the marker points which have at least one solid point near the boundary. The procedure for finding solid, fluid and marker particles near the boundary are shown in Fig. 2.

As mentioned earlier, all of the simulations were performed on the STL files of coral geometries, which were composed of millions of triangles. A ray-tracing algorithm was implemented to establish a relationship between these triangles with a Eulerian grid.The coordinates of the vertices and normal of each triangle are stored in the STL file. The ray triangle intersection algorithm used in the reconstruction of the flow field consists of the following steps.

**Fig 2.**
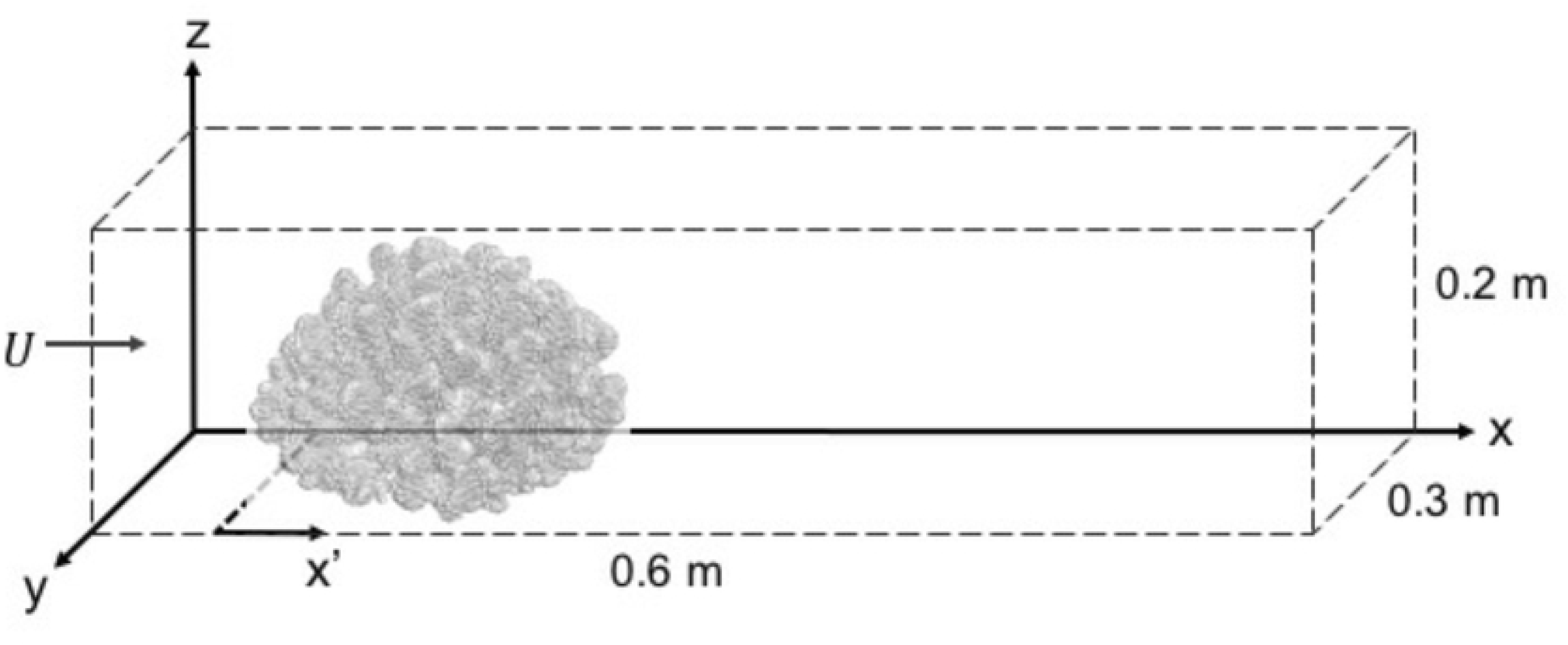
Tracking of solid-fluid interface. (A) An arbitrarily shaped solid on an orthogonal computational grid. The marker particles at the object’s boundary may or may not coincide with the Eulerian grid points. (B) Identification of solid and fluid points in the computational domain by ray tracing algorithm. (C) Identification of forcing points (where forcing will be applied) near the boundary after tagging solid and fluid grid points in the computational domain.

**Fig 3.**
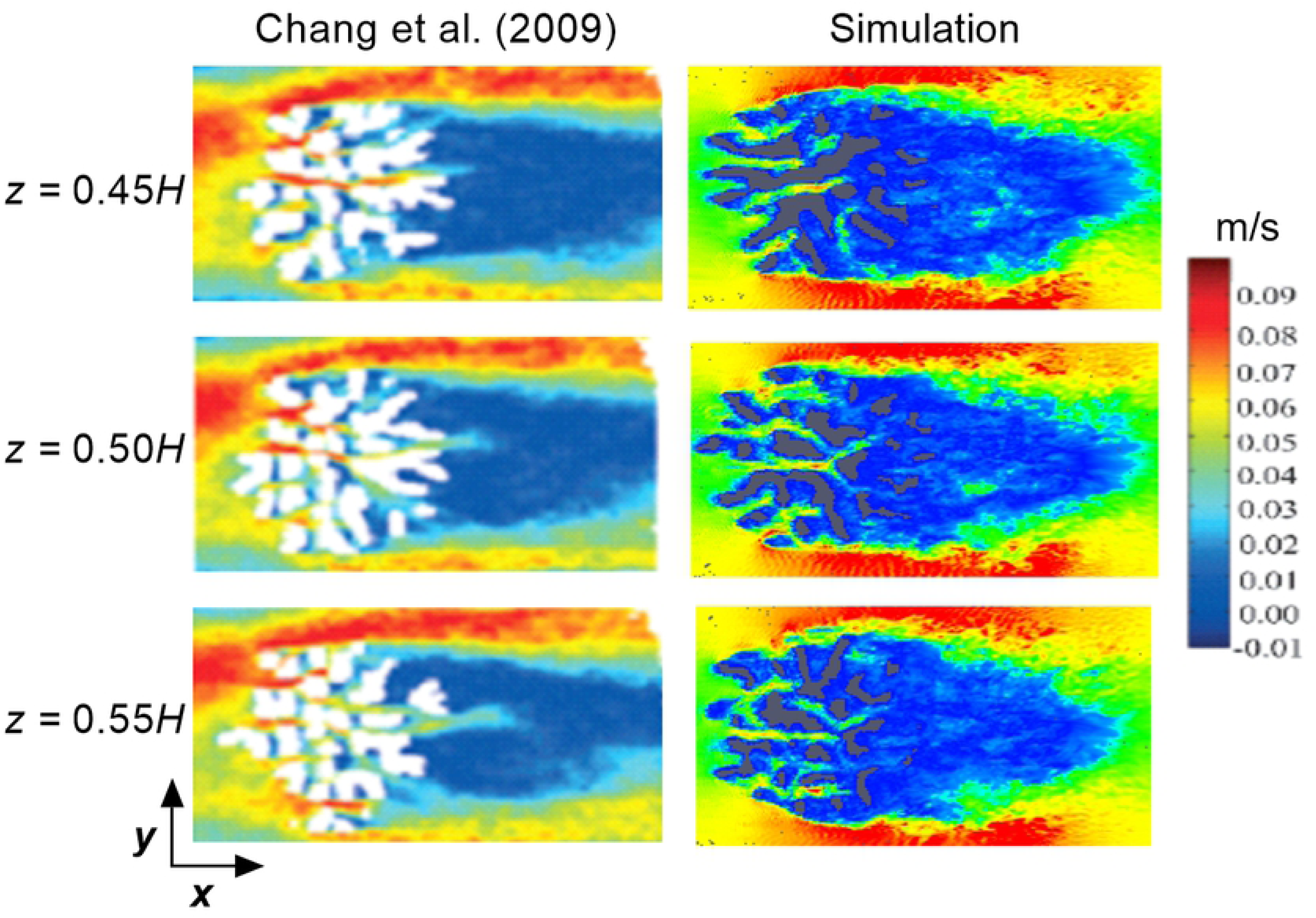
Schematic of the simulation flow domain (not to scale). *x′* = 0 is the secondary axis used at the beginning of the coral colony. The center of the coral is located at the middle of the flow domain. Uniform grid spacing is used throughout the domain.

1. A ray is shot from a grid point to a triangle.
2. Determine if the ray intersects the triangle or not.
3. Find the coordinates of the intersecting point.

In the solution procedure, the fluid points are the unknowns and the forcing points are boundary points, while the solid points do not influence the rest of the computation.

If the boundary coincides with the grid points, the forcing function can be calculated directly from the known 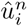 at the grid points. Otherwise, the value of velocity at the forcing points can be calculated from the interpolation of surrounding fluid points. The interpolation stencil in this method, however, involves a search procedure for suitable fluid points, and those points in the stencil may contain points that do not belong to the neighboring grid lines of the forcing point. The detailed procedure of identifying the immersed boundary, calculating the forcing function near the boundary, and the interpolation scheme can be found in Balaras *et al.* [46]. Here, the forcing function was calculated only at the boundary.

The simulation domain for the current study was not chosen randomly. Before performing the current analysis, a near-as-possible validation simulation was performed to compare the current result with the experimental result of Chang *et al.* [30] for a high-flow morphology coral, *Stylophora pistillata*. For the current analysis, the domain of simulation was 0.6 m long in the stream-wise direction (x), 0.3 m wide (y) and 0.3 m in vertical direction (z), similar to the flow domain used in the experimental and numerical study of Chang *et al.* [30]. As in natural conditions, a single coral sits in the open flow domain or in a coral reef of multiple colonies and the natural flow always takes the path of least resistance. But, the flow domain chosen in the experiment and current simulation, which acts to constrain the flow field, will only be approximated in nature for a few rare coral colonies. If the simulation can be performed in an extremely large domain, that would represent the ideal flow scenario for a coral sitting in an open flow environment. At the same time, we have to be realistic regarding the computational expenses when performing a simulation around an extremely complex and arbitrary shape geometry. In addition, we want to stress the objectives and scale of the current study. We intended to analyze the detailed flow field through and around a single coral colony, where the length scale varies from an individual colony branch diameter to the length or diameter of a single colony, and the large-eddy simulation of the Navier-Stokes equations were used here to resolve the hydrodynamics of coral interior at these scales. These constraints have led us to limit the dimension of the simulation domain for a single coral. As the current analysis was not intended to resolve diffusion boundary layers less than 1 mm, which would be extremely difficult and computationally expensive for such arbitrary geometry, the current grid resolution of 1 mm in x, y and z directions was sufficient to meet the objectives of the current study. Similar to Chang *et al.* [36], Dirichlet boundary conditions were applied at the inlet, and an outflow boundary was used at the outlet of the domain. For the lateral direction, slip boundary conditions were implemented at both sides of the coral. Slip boundary conditions were also used at the top face of the domain, and the bottom of the domain was defined as a no-slip wall. In the current analysis, 54 million grid cells were used for all of the simulations. All of the numerical experiments were performed on the Engineering Science and Mechanics (ESM) computing cluster at Virginia Tech. The ESM Linux Computational Cluster (LCC) consists of up to 52 compute nodes with 776 processors, 2TB RAM, and FDR-10 RDMA fabric connecting all of the nodes’ memory. Each of the nodes contains 20 processor cores and 24GB RAM.

## Results and discussion

Before comparing flow profiles between two *Pocillopora* structures, the results from the simulation were compared with the experimental results of Chang *et al.* [30] for the high-flow morphology of *Stylophora pistillata* for comparison. Although every coral is unique in nature, the dimensions of the corals used in the experiment and simulation were within the same order. In the experiment, the stream-wise (x), cross-stream (y), and vertical (z) extents of the densely-branched coral were 0.128 m, 0.13 2 m and 0.09 m, respectively. In the simulation, the extents of the densely-branched coral were 0.172 m, 0.172 m and 0.111 m, respectively, in the stream-wise, lateral and vertical directions. The channel dimensions for both the experiment and simulation domains were almost the same. In the experiment, the bulk velocity in the channel was 0.05 m/s and the Reynolds number, based on the diameter of the branch, was 580. Based on the height of the coral, it was approximately 5,000. Similar flow parameters were used in the simulation to keep the flow conditions close to those of the original experiment. Fig 4 shows a comparison of the cross-sectional (top view) velocity profile obtained near the midsection of the coral between the current simulation (right) and the experimental results of Chang *et al.* (left) at the different heights inside the corals. For the experiment, the middle slice was obtained exactly at the mid-height of the coral and the slices were obtained at 5-mm intervals from the both sides. For the simulation, the slices were chosen in exactly the same manner. For the simulation, high velocity was observed at both sides of the coral, and a distinct wake region formed behind the branches of coral. Because of the dense branch structure, most of the flow was diverted outward and vertically upward. The branches located in the frontal half of the coral significantly reduced the velocity for the lower half. The computational results were compared with the velocity slices obtained from the experiment. Although the oncoming velocity profile in Chang *et al.* is somewhat skewed owing to a curved, recirculating water channel, both velocity fields demonstrate the same characteristics for these flow conditions. The colony largely acts as a baffle, and high velocity regions, or ”jets,” whose speeds exceed the oncoming flow speed, can be found deep in the coral interior. The results also demonstrated a low velocity region behind the thick branches of the coral, and the velocity magnitude was almost identical for both cases.

**Fig 4.**
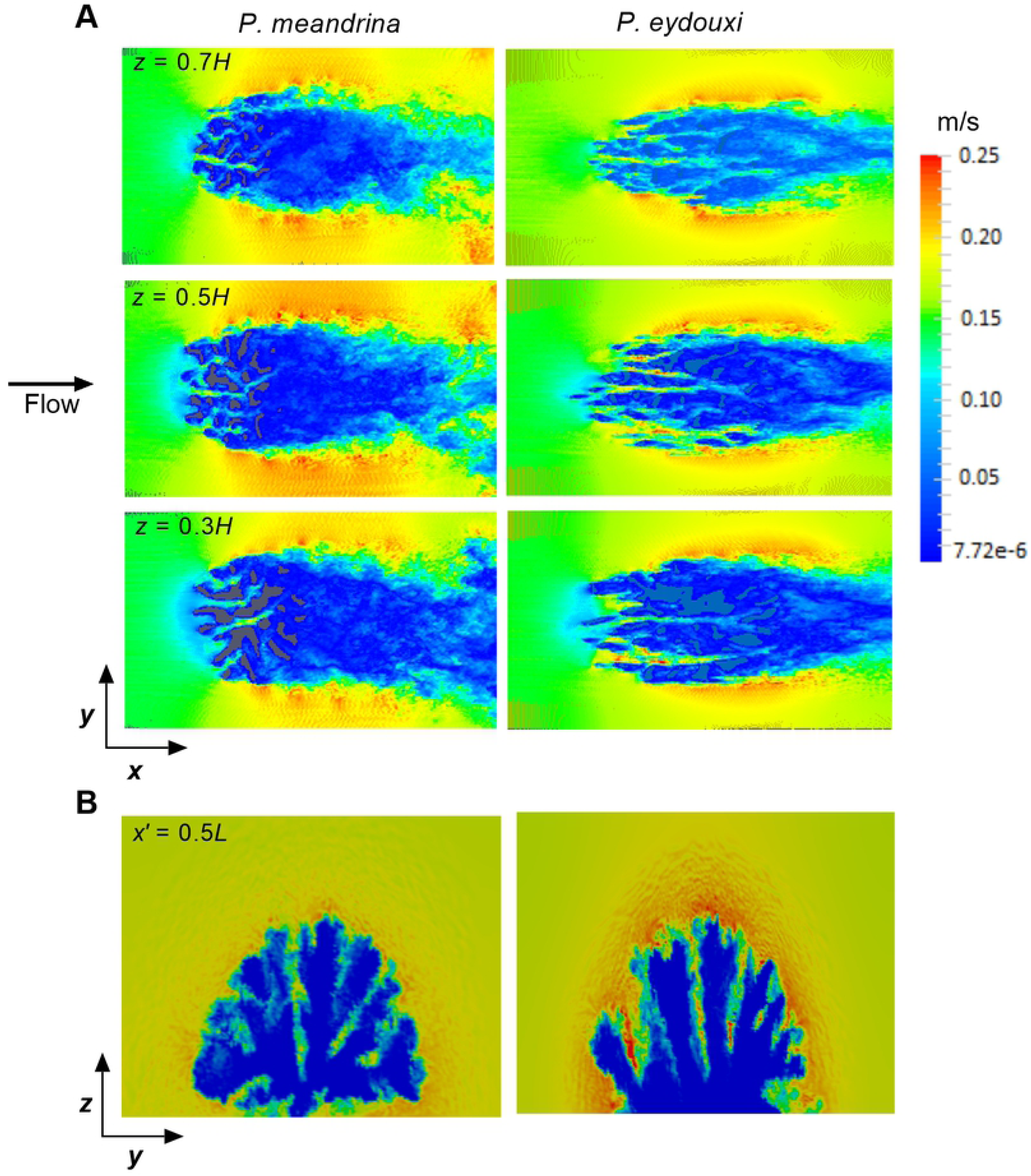
Numerical validation. Comparison of top view velocity magnitude plots between experimental results [30] and the present simulation at different heights in two coral colonies. On the left are velocity fields inside an *S. pistillata* colony obtained by magnetic resonance velocimetry at the approximate mid-height of the coral. (Reproduced with permission from *Limnology and Oceanography* 54, 1819 (2009). Copyright 2009 by John Wiley and Sons, Inc.) On the right are plots of the velocity field magnitude from the present LES simulations of flow through a *P. meandrina* colony, at the same heights.

Flow inside the coral is a function of inter-branch distance, branch diameter, coral height, and incoming velocity. To understand the branching effect of coral on the surrounding fluid, flow through two different branching patterns of the same *Pocillopora* coral was evaluated and compared for the same incoming flow. Top views of flow through these two *P. meadrina* corals were shown in Fig 5 for an incoming flow of 0.15 m/s. The incoming velocity was selected based on the high range of velocity magnitude near the *Pocillopora* coral habitat. The middle velocity slice was obtained at approximately the midsection of the coral and the rest of the slices were obtained at 20-mm intervals on both sides. For both of the coral, a stagnation region formed near the front section, and the effect was higher for *P. meandrina* due to high branch density. For *P. meandrina*, higher velocity was observed at the top and both sides of the coral and the velocity reduced substantially and became almost uniform in the latter half of the coral. As most of the flow is diverted to the top and both sides of the coral, the restriction of flow due to the high compactness of the interior branches was more visible for *P. meandrina*. During interaction with internal branches, mixing and transportation takes place at the coral surface. At the downstream end of the coral, a distinct wake region containing a large circulating eddy formed behind the coral. In contrast, the flow penetration at the interior of *P. eydouxi* was easier than in *P. meandrina*. As the geometry became more open, the interior received more flow, which was apparent from the flow slices exhibiting higher velocity zones at the interior of the coral. Though a large portion of the flow was diverted outward, the penetration was much better in *P. eydouxi* than *P. meandrina*.

**Fig 5.**
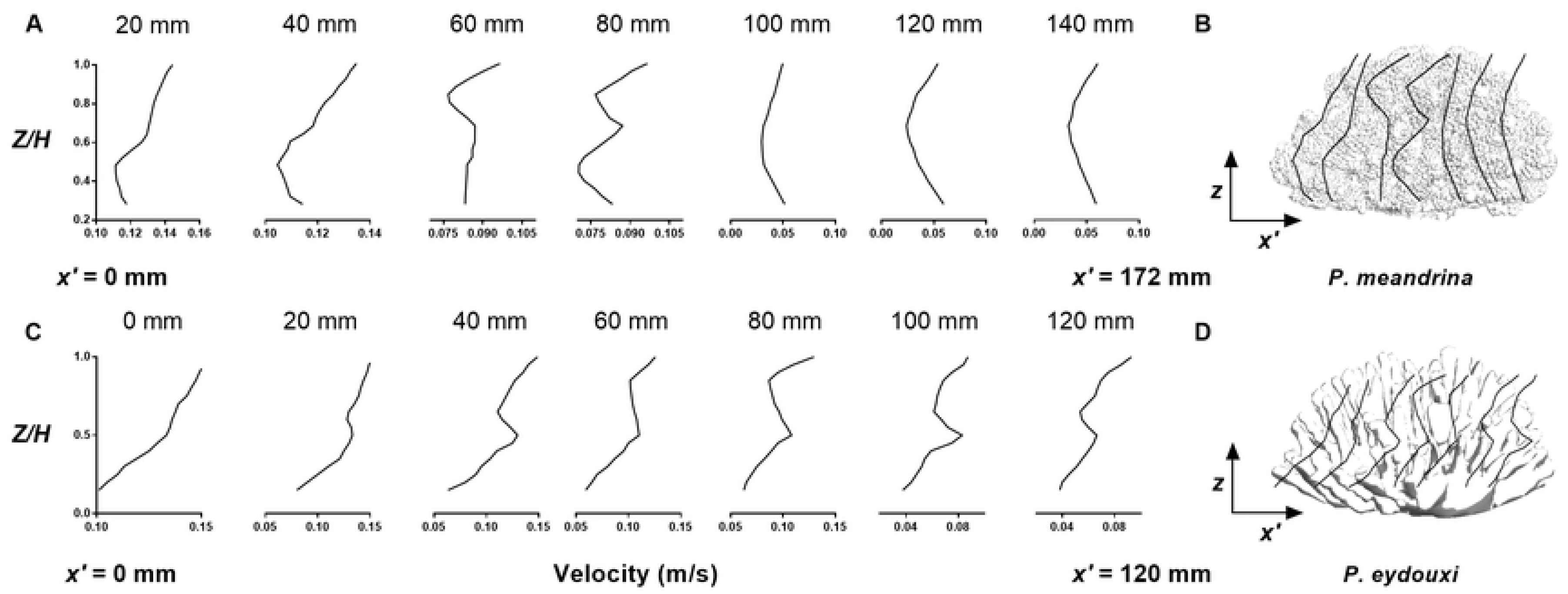
Comparison of two-dimensional velocity fields in *Pocillopora* colonies with different branch densities. (A) Top view velocity magnitude plots inside *Pocillopora meandrina* (left, relatively densely branched) and *Pocillopora eydouxi* (right, relatively loosely branched) colonies at *z* = 0.3*H*, 0.5*H* and 0.7*H* from the base of the coral for the same incoming velocity of 0.15 m/s. (B) Comparison of velocity magnitude plots in a plane perpendicular to the flow direction between *P. meandrina* (left) and *P. eydouxi* (right) at *x′* = 0.5*L*, the mid-length of the coral in the streamwise direction.

For qualitative measurements of the variation of velocity profiles along the lateral direction, a velocity slice (perpendicular to flow) was taken at *x′* = 0.5*L* for both of the corals. Fig 5 (B) shows a comparison of velocity slices obtained at the middle of the coral, perpendicular to the flow direction. For both of the corals, the outer periphery shows higher velocity magnitudes due to the constraints of the flow domain. For the same incoming flow conditions, *P. meandrina* shows relatively low velocity magnitude at the coral interior and the incoming flow loses most of its momentum due to high branch density. In comparison, *P. eydouxi* shows relatively higher velocity magnitudes between the branches, especially at the narrow corner between the branches, where the flow accelerates.

Up to now, we have provided qualitative comparisons of flow between these *Pocillopora* structures. But for better comparison, we require quantitative measurements of flow between these two corals. To reach this objective, mean vertical velocity profiles were obtained inside both of the corals at 20-mm intervals along their length. As the branches are extended randomly, it is difficult to obtain mean velocity profiles at different sections inside the coral. To overcome this issue, vertical velocity profiles were calculated from the average velocity magnitudes of the closest 10 neighbor grids at each location in the *X − Z* planes, and then a second average was performed along the lateral direction (*y* axis) within the coral. These vertical velocity profiles not only depicted the progression of flow inside the coral, but also indicated the amount of mixing and transportation that takes place along the length of the coral. The mean vertical velocity profiles obtained for both the *Pocilloppra* corals are shown in Fig 6 (A) and (C). For *P. meandrina*, the mean vertical velocity profiles were almost the same in nature up to 40 mm. This was the frontal portion of the coral, where the flow penetration was easy and the maximum velocity dropped approximately 33% of incoming velocity at the interior. For 60 to 80 mm, the flow started to slow down and dropped to a maximum 50 % of the incoming flow. At this section, the velocity profiles changed substantially. Interestingly, at 80 mm, the profile displayed an increase in velocity magnitude, where we expected further decline in incoming velocity. The internal structure of coral might accelerate the flow inside the coral, which may be necessary for the survival of internal section of the coral. From 100 mm up to the end of coral, the velocity profiles were almost identical and displayed a reduction in velocity magnitude at mid-height of the coral. In general, the velocity profiles showed low velocity at the middle of the coral and comparatively higher velocity at the top and bottom sections of the coral. Therefore, the nutrient availability will be higher in those regions than in the middle portion of the coral.

**Fig 6.**
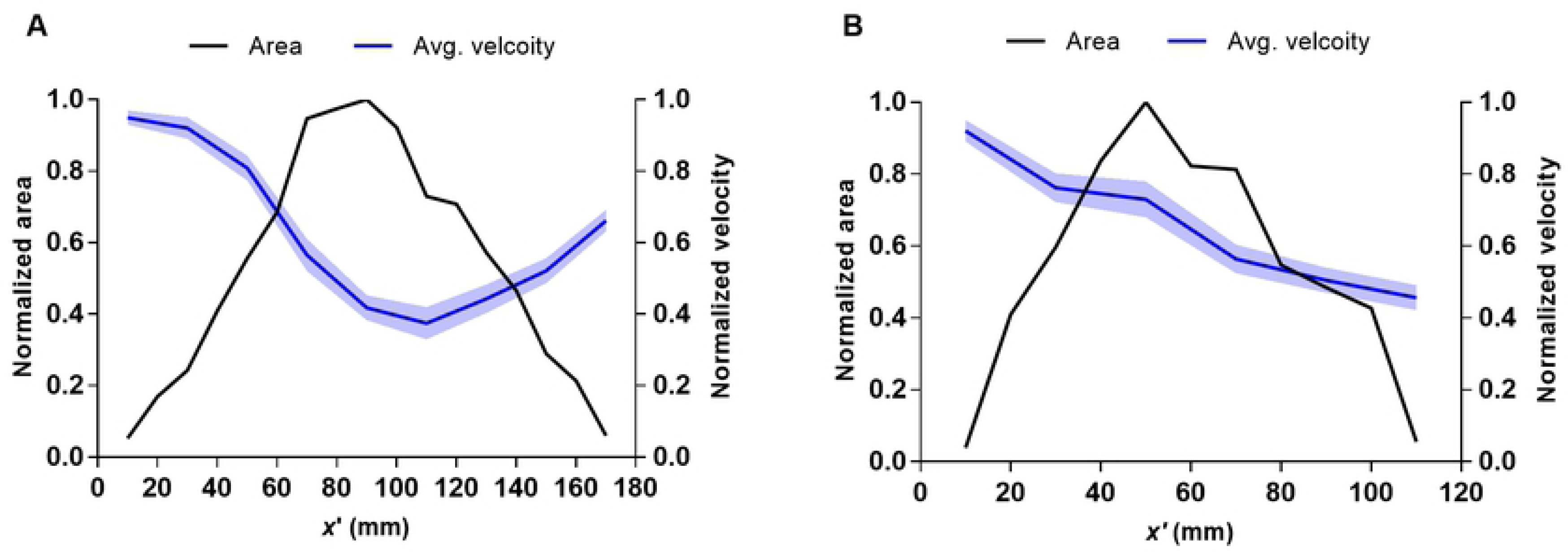
Variation of mean vertical velocity profiles in *Pocillopora* colonies with different branch densities. (A, C) Mean velocity profiles inside *P. meandrina* (A, relatively densely branched) and *P. eydouxi* (C, relatively loosely branched) were calculated at 20-mm intervals along the length of the coral by averaging the velocity magnitudes of the closest 10 neighboring grid points at several locations in the *x*-*z* planes, and then averaging these velocity magnitudes over the lateral (*y*) direction within the colonies. (B, D) Schematic of locations of the mean velocity profiles within the colonies (not to scale).

In contrast, for *P. eydouxi*, the velocity profile displayed higher values at the top of the coral, but lower values in the lower section. The vertical velocity profiles within *P. eydouxi* remained almost the same. Due to the openness of the structure, the flow did not vary much within the top half of the coral, where the flow magnitude was comparatively higher than in the lower half of the coral. For *P. eydouxi*, the velocity reduced abruptly at this lower section due to the solid hemispherical structure of coral at this section. At z = 0.5 H, the velocity magnitude increased due to the accelerated flow in the reduced area of the lower section of the coral. After that, the velocity dropped in the middle of the branches until the velocity regained its maximum magnitude, similar to that of the incoming flow at the top of the coral. Then, at the end section of coral (between 80 and 120 mm) the maximum velocity dropped to almost 50% of the incoming flow.

In addition to calculating vertical velocity profiles in different sections of these corals, mean velocity was also calculated along the length of two corals for the same incoming velocity. Fig 7 shows the variation of the mean velocity (averaged over the y-axis within the coral) with the cross-sectional area inside the coral. From the figure, it can be observed that the cross-sectional area was almost symmetrical for *P. meandrina* along the stream-wise direction. For densely-branched *P. meandrina*, the magnitude of incoming flow reduced significantly at the middle of the coral and recovered up to 60% at the end section. In contrast, the cross-sectional area was not symmetrical for *P. eydouxi* and the flow reduced to approximately 50% of the incoming flow along the length of the coral, which was lower than the flow in *P. meandrina* at the end of coral. But, within the coral, the overall magnitude of velocity was higher in *P. eydouxi* than in *P. meandrina* through the complete length of the coral. In terms of food, the interior of the coral has a strong chance of starving if the velocity is not sufficiently high. For *P. eydouxi*, the situation is comparatively better, as the interior of the coral receives a higher flow, which means better mass transport and food availability. However, the conditions for the coral that lives in the wake region are worse, because it takes a comparatively longer distance to recover flow in this area. This means that the nutrient availability for coral in the wake region will be reduced in comparison to the nutrient availability for coral living in the front region.

**Fig 7.**
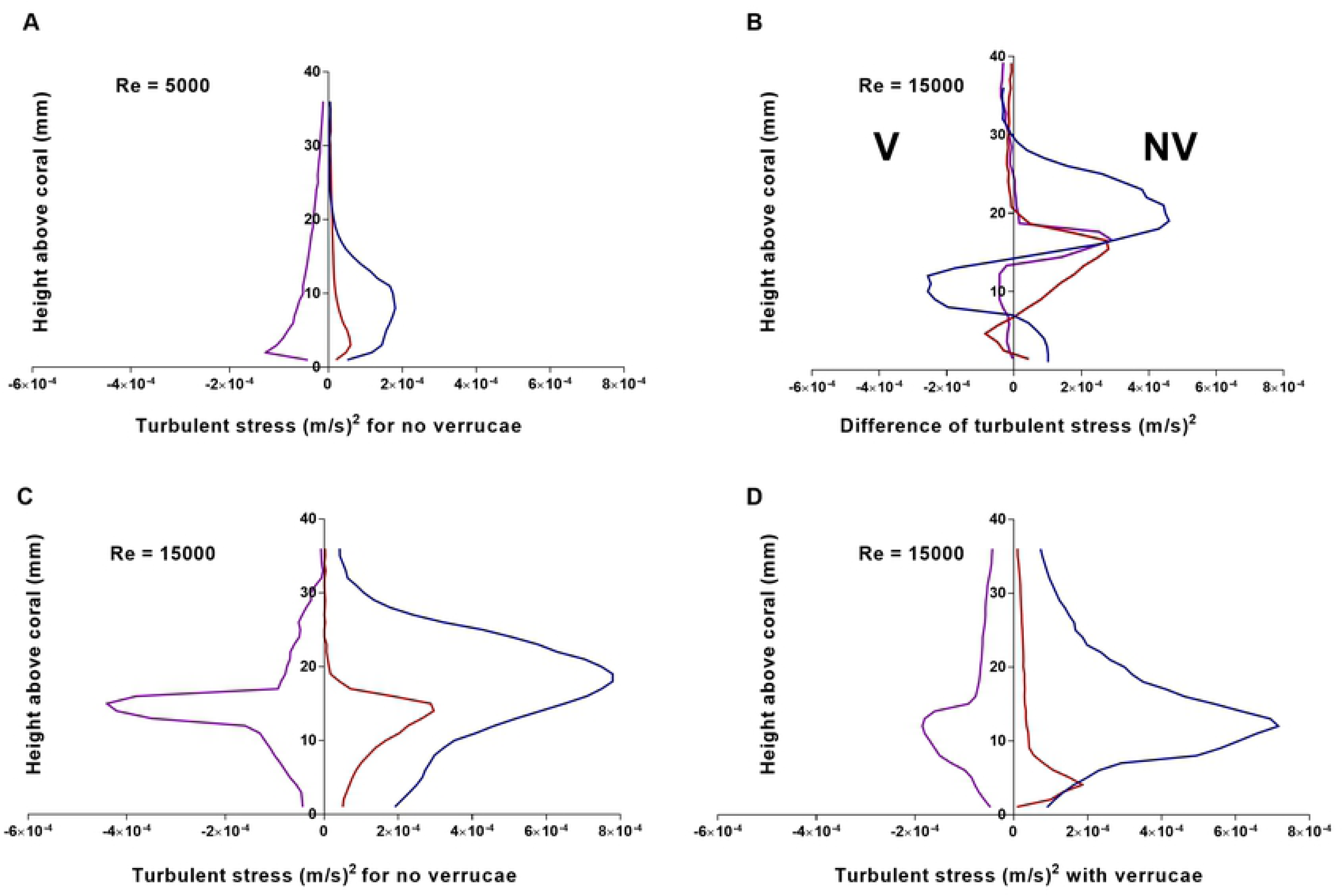
Variation of mean cross-sectional area and velocity in *Pocillopora* colonies with different branch densities. The normalized colony cross-sectional area and intra-colonial velocity values averaged over two-dimensional planes at eleven locations along the height of the colony for *P. meandrina* (A, relatively densely branched) and *P. eydouxi* (B, relatively loosely branched). Shading in the velocity variable represents one standard deviation from the mean (N=6). While the densely branched colony (A) displays minimum mean velocity values just above the mid-height of the colony, the minimum mean velocity values for the loosely branched colony (B) appear at the top of the colony.

### Comparison of flow dynamics over *M. capitata*

For a single coral, nutrient transfer and gas exchange depends on shear, turbulent mixing and the concentration gradient near the surface. Quantifying the turbulent stress is essential for explaining mixing, stress and drag developed on the coral surface. Even small changes in flow conditions can change these turbulent statistics over the boundary layer. As mentioned earlier, *M. capitata* with verrucae scattered on the coral surface grows near Kanehoe Bay in Hawaii where the flow condition is strong. To ascertain the impact of verrucae in natural flow conditions, computational analysis was performed on two different coral morphologies of loosely-branched *M. capitata* as described in the coral geometry section (Fig 8). To characterize the boundary layers and the turbulent stress developed on both of these structures, velocity profiles were computed over the coral surface at both weak and strong flow conditions. Based on the low-to-high water velocity near Kanehoe Bay and the height of the coral, Reynolds numbers 5,000 and 15,000 were used to simulate the weak and strong flow conditions for *M. capitata*. The comparison also highlights how the flow profiles change when the flow conditions vary from weak to strong for open coral structures. For ease of discussion, we have used CV and CWOV for the abbreviation of *Montipora capitata* with and without verrucae, respectively, hereafter. Fig 8 shows slices of the flow field around *M.Capitata* with and without verrucae at Reynolds number 15,000. After capturing flow fields for both the structures, velocity and stress were calculated at *x′* = 0.7*L* above the surface of the corals at two different Reynolds numbers. Then, the velocity and stress profiles obtained were averaged along the lateral (y) direction within coral to obtain a mean of the profiles on top of the coral’s surface. The mean profiles were compared for CV and CWOV at the same flow conditions to capture the effects of verrucae on the flow field clearly.

**Fig 8.**
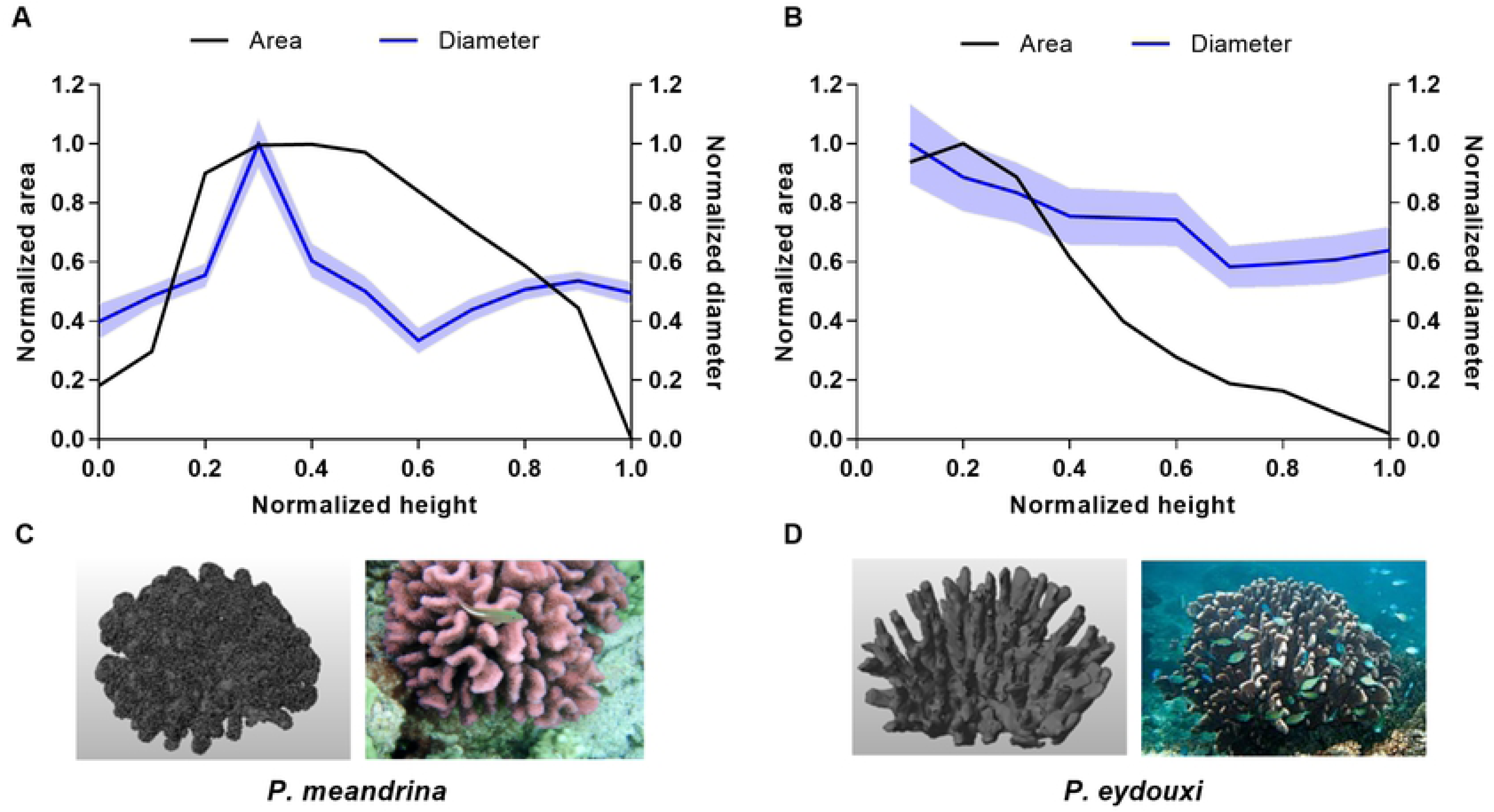
Comparison of two-dimensional velocity fields in a *Montipora* colony with and without roughness elements. (A) A *Montipora capitata* colony with the roughness elements (verrucae) removed, and (B) the same colony with the verrucae intact, at a Reynolds number of 15,000. The velocity field plots were obtained at *x′* = 0.7*L*.

Fig 9 (A & B) shows a comparison of horizontal velocity profiles at the top of *Montipora capitata* with and without verrucae at Reynolds numbers 5,000 and 15,000. For both cases, the mean velocity profile was normalized by the incoming velocity. For CWOV, the location of free-stream velocity was observed at about 9 mm above *Montipora capitata* at Reynolds number 5,000, whereas the distance to reach the maximum horizontal velocity magnitude doubled to 18 mm for CV. This demonstrated that the maximum velocity magnitude was found closer to the top of the coral surface at a higher Reynolds number. For CWOV, the inflection point was observed at about 5 mm above the coral surface at Reynolds number 15,00, and for CV, it was observed at approximately 11 mm above the surface.

**Fig 9.**
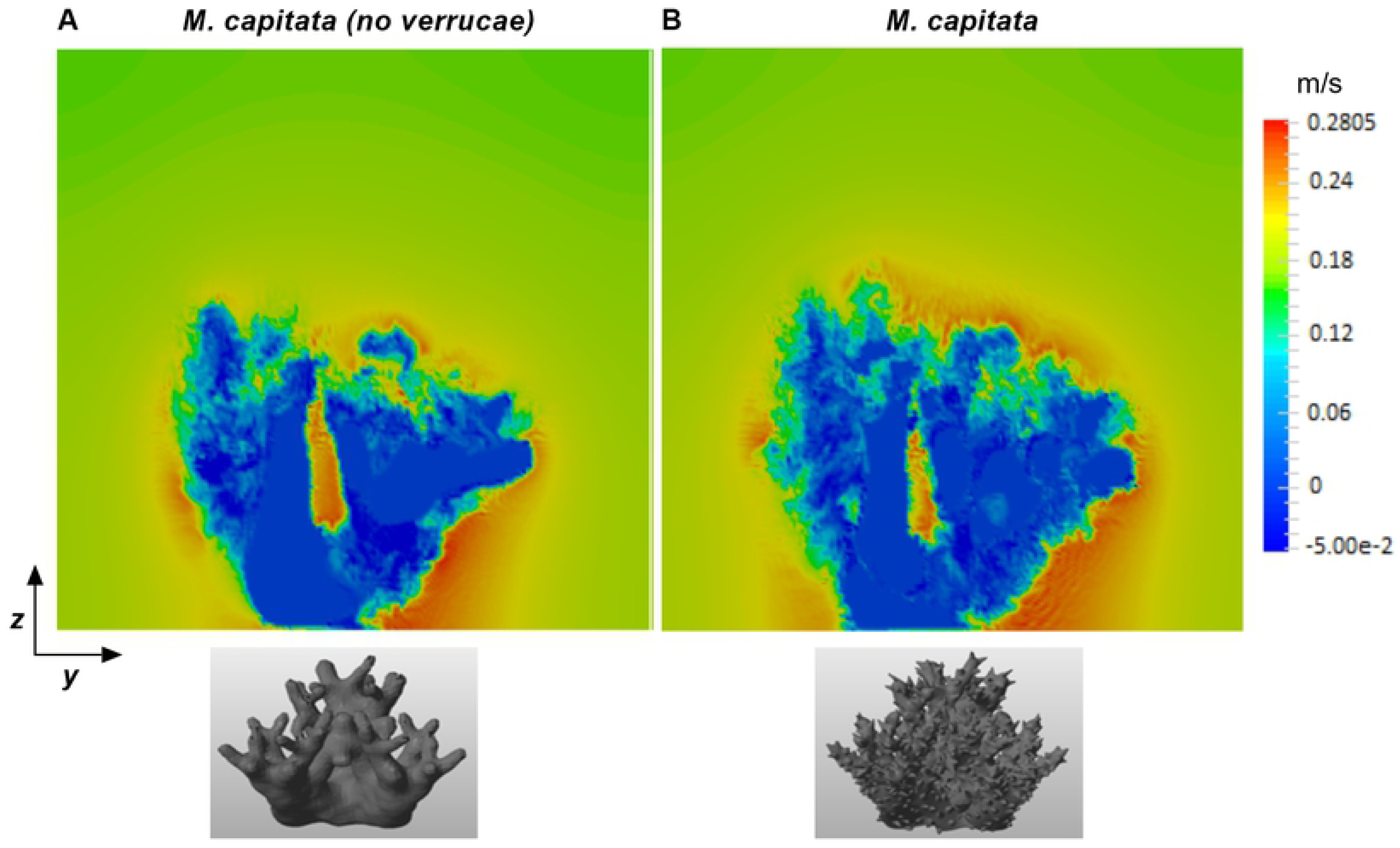
Comparison of velocity profiles above a *Montipora* colony with and without roughness elements. (A) The horizontal (*x*-direction) velocity profile at *Re* = 5, 000, and (B) the horizontal (*x*-direction) velocity profile at *Re* = 15, 000. (C) The vertical velocity profile at *Re* = 5, 000, and (D) The vertical velocity profile at *Re* = 15, 000. All profiles were obtained at *x′* = 0.7*L* and were averaged over the lateral (*y*) direction. The vertical axis indicates the height above the colony surface.

A comparison of the two horizontal velocity profiles at Reynolds number 5,000 reveals that the two profiles are almost identical at up to 75% of the normalized velocity. This indicates very similar flow profiles above the coral surface at 6 mm for CWOV and 7 mm for CV at this low Reynolds number. This means that the nutrient, or mass, transfer rate along the horizontal direction should be similar for the two cases up to this height. After this point, the transfer rate might change according to the velocity profile. In contrast, the two horizontal velocity profiles were different at Reynolds number 15,000. The two profiles reached 50 % of its maximum magnitude at about 2 and 5 mm above the surface of CWOV and CV respectively. At the same Reynolds number, the free-stream velocity was observed at approximately 5 mm above the surface of CWOV, whereas the free-stream velocity was found at approximately 9 mm above the surface of CV. If we continue to move vertically up the coral structures, the two velocity profiles collapsed at approximately 27 mm above the coral surface. In an experimental analysis, Reidenbach and Stocking *et al.* [41,47] calculated horizontal velocity profiles at the top of a single coral for unidirectional flow, and the velocity profile and the location of inflection points were almost the same in nature when compared to the velocity profiles computed in the current simulation.

As the coral depends on natural flow conditions for the transfer of nutrients and mass, the high velocity magnitude near the coral surface translates to fast mass and momentum transfer from the coral surface. This condition is very important for biological processes like photosynthesis and respiration. For photosynthesis and respiration, the gas produced near the surface should be removed quickly; otherwise, the gas produced on the surface may hinder the coral’s natural physiological processes. If the comparison is made between CWOV and CV, the mixing and transport should be better for CWOV at both of the Reynolds numbers, due to higher horizontal velocity magnitude near the coral’s surface. But, the higher velocity magnitude near the coral’s surface raises the question of surface integrity, which in turn brings the role of natural adaptability of biological systems to the fore. In strong flow conditions, the open-structure coral maintains a delicate balance between the fast mass transfer from the surface of the coral, which is required for nutrient transfer, and surface integrity, which is essential in protecting the branches of the coral.

Fig 9 (C & D) show the variation of vertical velocity profiles above the coral surface at Reynolds numbers 5,000 and 15,000 respectively. At Reynolds number 5,000, the two velocity profiles were almost the same for both cases, and the magnitude was smaller in comparison to the corresponding horizontal velocity. As the Reynolds number increased, the velocity profile for CWOV demonstrated a higher magnitude near the coral surface. For both cases, the maximum velocity region was found close to the coral surface, and the profiles reached a constant magnitude at approximately 15 mm above the coral surface.

If we compare the horizontal velocity to the vertical profile, the mass and momentum transport is mainly dominant in the horizontal direction at both of the Reynolds numbers. At Reynolds number 5,000, the two *M. capitata* corals exhibited similar kinds of transportation along the vertical direction, and the magnitude increased with increased Reynolds numbers. If the vertical flow profiles for these two coral structures are compared, CWOV should have better mixing and transportation along the vertical direction than CV. Also, the nutrient gradient above the canopy will be more uniform for CWOV due to comparatively higher vertical flow. In an experimental analysis completed by Reidenbach *et al.* [41]), the vertical velocity profiles measured above the coral surface displayed almost the same profile and magnitude for unidirectional flow.

#### Turbulent stress over coral surface

The transfer of nutrients and mass between coral and the overlying water column depends upon shear stress developed on the coral surface. To understand the effects of turbulent dynamics on the surface of coral, time-and-space averaged turbulent stress component <u′u′>, <w′w′> and <u′w′>, were computed for both of the coral surfaces for unidirectional flow where the shear Reynolds stress, <u′w′>, is mainly responsible for mixing near the coral surface. To capture the fluctuating quantities, the computation was given enough time for the stability of mean flow to become established. Profiles of Reynolds stress at the top of *Montipora capitata* without verrucae are shown in Fig 10 at Reynolds numbers 5,000 and 15,000.

**Fig 10.**
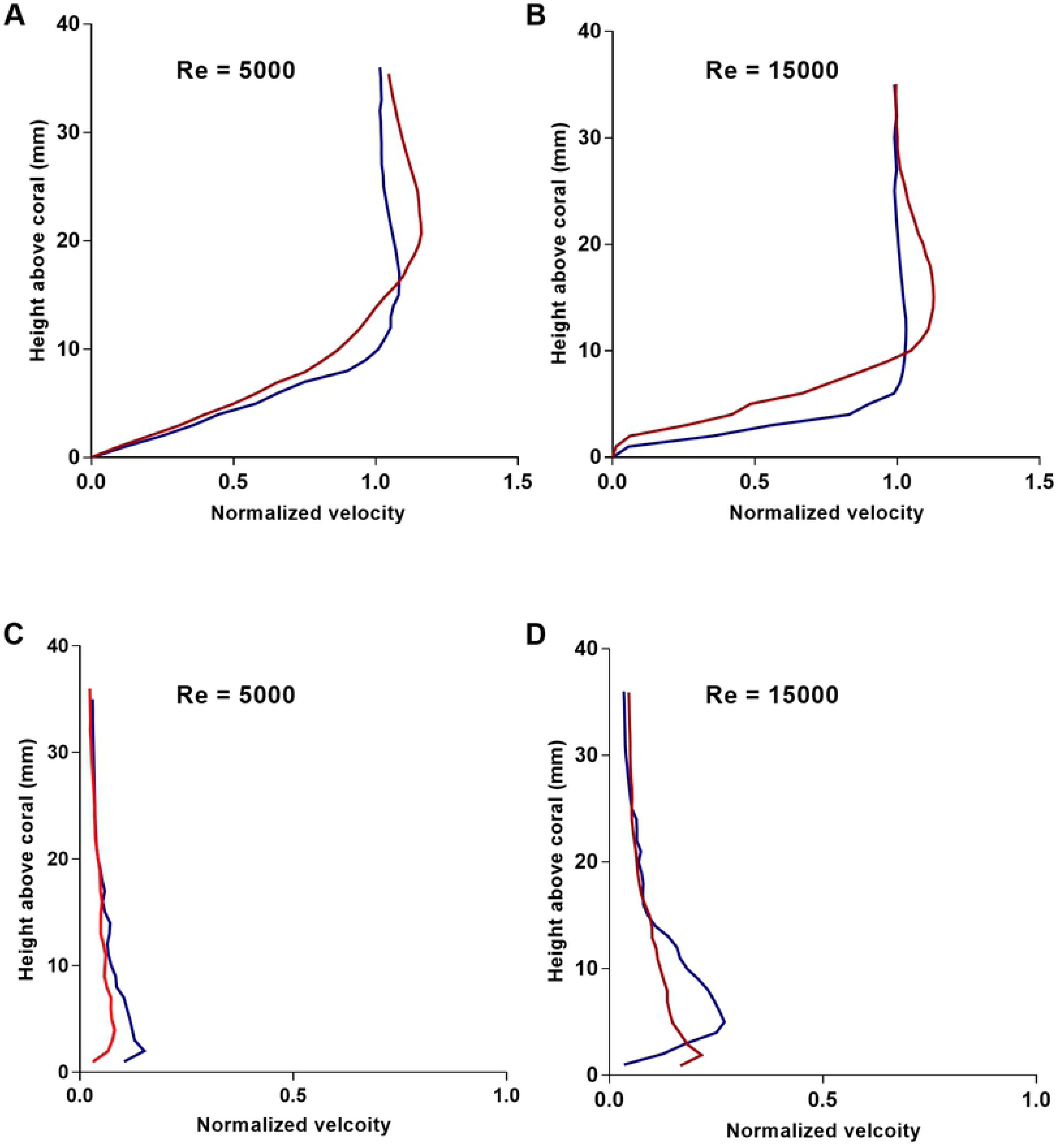
Comparison of turbulent stress components above a *Montipora* colony with and without roughness elements. (A) Turbulent stress components above a *Montipora capitata* colony without verrucae (roughness elements) at *Re* = 5, 000. (B) The difference between the turbulent stress components above the *Montipora capitata* colony with and without verrucae at *Re* = 15, 000. (C) Turbulent stress components above a *Montipora capitata* colony without verrucae at *Re* = 15, 000. (D) Turbulent stress components above a *Montipora capitata* colony with verrucae at *Re* = 15, 000. All profiles were obtained at *x′* = 0.7*L* and averaged over the lateral (y) direction.

At a Reynolds number of 5,000, the <u′u′> turbulent stress reached maximum magnitude at approximately 9 mm above the surface, indicating maximum horizontal momentum transport near the coral polyps. In contrast, both the <w′w′> and <u′w′> were smaller in magnitude. The existence of maximum Reynolds shear stress, <u′w′>, near the surface indicated maximum mixing at the surface of coral, and the magnitude approached zero at approximately 35 mm above the coral surface. The maximum magnitude of this cross value stress was almost 50% of the maximum value of <u′u′> stress, and several times higher than <w′w′>. When the Reynolds number increased from 5,000 to 15,000, the maximum magnitude of <u′u′> stress increased to almost four times the previous magnitude, with the maximum value found at approximately 19 mm above the coral surface, whereas <w′w′> and <u′w′> components reached maximum magnitude at around 15 mm. For <w′w′> stress, the magnitude came close to zero at around 20 mm above the coral surface for a Reynolds number of 15,000, whereas the maximum <w′w′> value was achieved at around 3 mm above the coral surface for Reynolds number 5,000. Overall, the results revealed a small magnitude of stress at Reynolds number 5,000, although the cross-value stress, <u′w′>, showed a comparatively higher value, even at such a low Reynolds number. This shows that mixing will be better for the smooth surface of CWOV even at Reynolds number 5,000. As the Reynolds number increased to 15,000, the magnitude of all of the Reynolds stresses increased and the mixing was expected to be better due to the contribution from the <u′w′> component.

Fig 10 (D) shows the components of Reynolds stress developed on top of *M. capitata* with verrucae at Reynolds number 15,000. Here, the maximum value for <u′u′> stress was found just below 15 mm above the coral surface. Though the maximum magnitude was almost the same as that of CWOV, the overall magnitude was higher for CWOV than for CV. For the vertical stress component, <w′w′>, the maximum value was found just about 5 mm above the coral surface for CV and the stress dampened very quickly. In contrast, the vertical stress dampened at approximately 20 mm above the coral surface for CWOV. This means that between both corals, CWOV had higher vertical transport than CV at the same Reynolds numbers. For both cases, the <u′w′> component approached zero at about 25 mm above the coral surface and remained almost constant for both structures. But below this height, the maximum magnitude of <u′w′> Reynolds shear stress for CV was almost twice that of CWOV. To validate these results, they were compared with the turbulent stress analysis on top of a single coral from the experimental study of Stocking *et al.* [47]. In the experiment, the Reynolds stress was measured as the covariance of the horizontal and vertical velocity fluctuations. The experimental analysis shows that the maximum magnitude of turbulent stress takes place within 20 mm above the top of the coral. In the current analysis, the locations of maximum stress for all of the components are found within 20 mm above the coral surface, and the magnitude of the stress components are within the same order of magnitude as those found in the experiment. In the experiment, the vertical stress, <w′w′>, was almost half of the horizontal stress, <u′u′>, but the simulation results show a comparatively lower value for vertical stress except for the CWOV at Reynolds number 15,000. Fig 10 (B) shows the difference of magnitude in turbulent stress developed at the top of two *M. capitata* at Reynolds number 15,000. In general, Reynolds stresses follow the typical boundary layer structure followed by the reef system.

## Conclusion

Computational analyses were performed to understand the variation of flow profiles inside two *Pocillopora* coral geometries with different branching patterns. Slices of flow profiles at the interior of the corals depicted the interaction of flow with coral branches at different sections of the coral. To quantify the variation, mean velocity profiles were estimated along the length of the coral at the interior. The mean vertical profile displayed higher velocity magnitudes at the top and base of *Pocillopora meandrina*. The dense branching pattern presented higher resistance to flow in the mid-section of the coral. The mean normalized velocity along the length of the coral dropped to approximately 38% in the middle of the coral, but recovered to 65% at the back of the coral. In contrast, the results displayed higher flow penetration in *Pocillopora eydouxi* through its sparser branches up to the mid-section, but the velocity dropped to approximately 50 % of the incoming flow. However, the mean velocity magnitude was higher than that of *Pocillopora meandrina* at the front and middle of the coral. If the comparison is made between the mean velocity profiles at the different sections of the coral, it becomes clear that the mean profile changes substantially at the middle of the coral for densely branched coral, whereas the mean profile remains almost the same for open structures all along the length of the coral. The comparison of flow for both of the structures also provides an indication of mass transfer and growth rates at different sections of the coral.

To understand the adaptability of coral, simulations were performed on *Montipora capitata* with and without verrucae at two different Reynolds numbers. The results displayed distinct inflection points and differences in velocity profiles for coral with and without verrucae for horizontal profiles at these Reynolds numbers. At both of the Reynolds numbers, the thickness of the horizontal velocity profiles for coral with verrucae were higher than for coral without verrucae. From the comparison between CV and CWOV, it can be concluded that the transport will be better in the horizontal and vertical directions for CWOV at both of the Reynolds numbers, due to higher velocity magnitudes near the surface of the coral. In addition to velocity profile, turbulent stress was also calculated for both morphologies. The results showed that the magnitude of the turbulent stress increased with the Reynolds number. At Reynolds number 5,000, the results showed a maximum magnitude for <u′u′> stress and comparatively higher Reynolds shear stress <u′w′> for CWOV, which indicates better mixing even at such moderate flow conditions. At Reynolds number 15,000, the magnitude for <u′u′> stress displayed almost the same magnitude for both of the geometries, but the vertical turbulent stress, <w′w′>, decreased in magnitude and dampened quickly for CV. In addition, the shear Reynolds stress, <u′w′>, demonstrated the maximum magnitude for CWOV over CV out of the two geometries. From the comparison of velocity and turbulent stress for both morphologies, it can be concluded that mixing and transportation will be better for CWOV over CV at the same flow conditions, but in terms of surface integrity, CV has the advantage over CWOV.

## Acknowledgments

We would like to thank Elias Balaras of George Washington University for the use of his immersed boundary code. We also want to thank Uri Shavit of Israel Institute of Technology (Technion) for providing the *P. meandrina* geometry file.

